# Coronavirus Resistance Database (CoV-RDB): SARS-CoV-2 susceptibility to monoclonal antibodies, convalescent plasma, and plasma from vaccinated persons

**DOI:** 10.1101/2021.11.24.469823

**Authors:** Philip L. Tzou, Kaiming Tao, Sergei L. Kosakovsky Pond, Robert W. Shafer

**Affiliations:** Division of Infectious Diseases, Stanford University School of Medicine, Stanford, CA, U.S.A.; Institute for Genomics and Evolutionary Medicine, Temple University, Philadelphia, PA, U.S.A.

## Abstract

As novel SARS-CoV-2 variants with different patterns of spike mutations have emerged, the susceptibility of these variants to neutralization by antibodies has been rapidly assessed. However, neutralization data are generated using different approaches and are scattered across different publications making it difficult for these data to be located and synthesized. The Stanford Coronavirus Resistance Database (CoV-RDB; https://covdb.stanford.edu) is designed to house comprehensively curated published data on the neutralizing susceptibility of SARS-CoV-2 variants and spike mutations to monoclonal antibodies (mAbs), convalescent plasma (CP), and vaccinee plasma (VP). As of October 2021, CoV-RDB contains 186 publications including 64 (34%) containing 7,328 neutralizing mAb susceptibility results, 96 (52%) containing 11,390 neutralizing CP susceptibility results, and 125 (68%) containing 20,872 neutralizing VP results. The database also records which spike mutations are selected during *in vitro* passage of SARS-CoV-2 in the presence of mAbs and which emerge in persons receiving mAbs as treatment. The CoV-RDB interface interactively displays neutralizing susceptibility data at different levels of granularity by filtering and/or aggregating query results according to one or more experimental conditions. The CoV-RDB website provides a companion sequence analysis program that outputs information about mutations present in a submitted sequence and that also assists users in determining the appropriate mutation-detection thresholds for identifying non-consensus amino acids. The most recent data underlying the CoV-RDB can be downloaded in its entirety from a Github repository in a documented machine-readable format.

## Introduction

Beginning in late 2020, several SARS-CoV-2 variants sharing multiple spike mutations were reported from different parts of the world. These variants have been classified according to their phylogenetic lineage and component mutations. Variants that spread widely and displayed evidence for being more transmissible, causing more severe disease and/or reducing neutralization by antibodies generated during previous infection or vaccination have been classified as variants of concern (VOCs) by the World Health Organization and U.S. Centers for Disease Control and Prevention (reviewed in [1]). Variants that have spread less widely but share some of the key mutations present within VOCs have been classified as variants of interest (VOIs).

As novel SARS-CoV-2 variants have emerged, numerous investigations assessed the susceptibility of individual and combination spike mutations to neutralization by monoclonal antibodies (mAbs), convalescent plasma (CP), and vaccinee plasma (VP). The susceptibility of SARS-CoV-2 variants to mAbs is obviously relevant for preventing and treating SARS-CoV-2 infections with mAb regimens. The susceptibility of SARS-CoV-2 variants to CP provides insight into the likelihood that a variant will infect and cause illness in a person previously infected with and recovered from a different variant. The susceptibility of SARS-CoV-2 variants to VP provides insight into the risk that a variant will infect and cause illness in a previously vaccinated person.

The SARS-CoV-2 spike protein is a 1,273-amino acid trimeric glycoprotein responsible for entry into host cells. Each spike monomer has a largely exposed S1 attachment domain (residues 1–686) and a partially buried S2 fusion domain (residues 687–1,273) [2, 3]. Part of S1, called the receptor-binding domain (RBD; residues 306–534) binds to the human angiotensin-converting enzyme 2 (ACE2) receptor [4, 5]. Approximately 20 RBD residues bind the ACE2 receptor. The part of the RBD containing these residues (438-506) is referred to as the receptor-binding motif (RBM), whereas the remainder of the RBD is called the RBD core. The SARS-CoV-2 RBD is the main target of neutralizing antibodies [6, 7]. Like the RBD, much of the S1 amino-terminal domain (NTD) is also exposed on the spike trimer surface and is targeted by neutralizing antibodies.

The Stanford Coronavirus Resistance Database (CoV-RDB) provides open access to continuously curated neutralizing susceptibility data of SARS-CoV-2 mutations and variants to mAbs, CP, and VP. The database includes specific data on which spike mutations have been selected during both i*n vitro* passage of SARS-CoV-2 in the presence of mAbs and *in vivo* emergence in persons receiving mAbs for treatment. The CoV-RDB website (https://covdb.stanford.edu) enables users to query its database and the associated Github repository enables users to download the entire database. The CoV-RDB website also contains a companion sequence analysis program that annotates SARS-CoV-2 user-submitted sequences using CoV-RDB data.

## Methods and results

The CoV-RDB data model is made up of four major entities: published references, viral mutations and variants, antibodies (including mAbs, CP, and VP), and experimental results. As of October 2021, the database contains 186 publications: 18 (10%) with 380 results from in vitro mAb selection experiments, 64 (34%) with 7,328 neutralizing mAb susceptibility results, 96 (52%) containing 11,390 with CP susceptibility results, and 125 (68%) with 20,872 neutralizing VP results. As publications often contain more than one type of experiment, the sum of the percentages is greater than 100%. The complete database schema is available at https://github.com/hivdb/covid-drdb/blob/master/schema.dbml.

### Curation of published references

Published references included in the CoV-RDB are obtained weekly from three literature sources:i) PubMed using the search term “SARS-CoV-2”; ii) BioRxiv/MedRxiv COVID-19 SARS-CoV-2 preprint servers; and iii) the Research Square SARS-CoV-2/COVID-19 preprint server. Publication titles and abstracts are reviewed manually to identify studies containing data on SARS-CoV-2 spike mutations and humoral immunity. Studies that pass initial review are downloaded to a Zotero reference database and full texts are reviewed to extract specific data on the selection of spike mutations *in vitro* and *in vivo*, and neutralizing susceptibility data for individual spike mutations or combinations of spike mutations, such as those present in SARS-CoV-2 variants to mAbs, CP, and VP. Curated studies initially published as preprints are re-evaluated following peer-review publication.

For each publication, data are first entered into linked comma-separated files (CSVs) contained in a Github repository (https://github.com/hivdb/covid-drdb-payload). Data are then imported into a Postgres database where they are validated for completeness and consistency. The data in the Postgres database are then exported as a single SQLite database file that is both available for download under the CC BY-SA 4.0 open-source license and used as the source for regularly updated datasets and queries on the CoV-RDB website. The entire workflow is summarized in S1 Fig.

### Virus variants and mutations

In the CoV-RDB data model, the virus entity represents individual spike mutations, combinations of spike mutations, and virus variants for which the full set of spike and genomic mutations is known including VOCs and VOIs. Published *in vitro* neutralization experiments have been performed using (i) pseudotyped viruses (vesicular stomatitis viruses [VSV] and lentiviruses) containing a SARS-CoV-2 spike protein with specific mutations; (ii) chimeric SARS-CoV-2 viruses in a VSV genomic backbone containing a spike protein with specific mutations; (iii) full-length cloned SARS-CoV-2 recombinant viruses generated using multiple plasmids, or bacterial or yeast artificial chromosomes; and (iv) primary SARS-CoV-2 isolates [1]. For pseudotyped, chimeric, and recombinant viruses, the virus can be characterized entirely by its spike mutations. For the primary SARS-CoV-2 isolates, the virus is also characterized by mutations in other viral proteins that may influence viral replication kinetics but not neutralization susceptibility. Most *in vitro* selection experiments have been performed using chimeric VSVs containing SARS-CoV-2 spike proteins as these viral constructs can undergo multiple rounds of replication in cell culture.

Table 1 lists the VOCs, VOIs, and other full-genomic SARS-CoV-2 variants for which data are available in CoV-RDB. Approximately 85% of results on variants are for VOCs, 12% are for VOIs, and 3% are for other variants containing one or more RBD mutations. As SARS-CoV-2 variants are continually evolving and variant definitions are regularly updated, individual VOCs and VOIs often differ slightly in the mutations that they contain. S1 Table displays the variability in these patterns for each of the VOCs and VOIs. For example, the alpha, beta, gamma, and delta variants in CoV-RDB contain 11, 20, 7, and 17 unique patterns of spike mutations, respectively, all closely related to an archetypal consensus variant.

**Table 1.** Variants of Concern, Variants of Interest, and Other Variants for which Neutralization Data are Available in CoV-RDB.

| **Virus types** | **References** | **Results** |
| --- | --- | --- |
| ***Variants of concern*** |  |  |
| Alpha | 106 | 7439 |
| Beta | 114 | 7823 |
| Gamma | 61 | 3247 |
| Delta | 40 | 2612 |
| ***Variants of interest*** |  |  |
| Epsilon | 22 | 926 |
| Eta | 9 | 216 |
| Iota | 14 | 554 |
| Kappa | 23 | 823 |
| Lambda | 8 | 324 |
| Mu | 4 | 32 |
| ***Other variants containing $\geq 1$ RBD mutation (n=17)*** |  |  |
| B.1.463 [F384L] | 1 | 220 |
| B.1.1.298 [Y453F] | 6 | 192 |
| B.1.617.3 [L452R, E484Q] | 2 | 100 |
| B.1.1.241 [S477N] | 1 | 64 |
| B.1/E484K | 1 | 32 |
| R.1 [E484K] | 1 | 32 |
| B.1.1.141 [T385I] | 1 | 17 |
| B.1.1.317 [S477N, A522S] | 1 | 14 |
| B.1.1.298 [Y453F] | 6 | 192 |
| B.1.1.318 [E484K] | 1 | 25 |
| B.1.618 [E484K] | 1 | 20 |
| A.23.1 [V367F, E484K] | 3 | 27 |
| B.1.1.519 [T478K] | 2 | 12 |
| A.30 [R346K, T478R, E484K] | 1 | 8 |
| A.27/A222V [L452R, N501Y] | 1 | 6 |
| B.1.619 [N440K, E484K] | 1 | 1 |
| R.2 [E484K] | 1 | 1 |
| **Footnote:** The most common ancestral variants lacking immune escape mutations used as controls include A variants (n=3899), B variants (Wuhan consensus; n=17136), B.1 variants (Wuhan consensus + D614G; n=16387). |  |  |

Table 2 lists the most common studied individual mutations within the spike RBD, NTD, C-terminal domain (CTD), and S2 domain. These mutations have generally been studied within an ancestral virus backbone. CoV-RDB contains experimental data on 296 different individual spike mutations at 159 positions including 197 RBD mutations at 76 positions, 55 NTD mutations at 43 positions, 32 S2 mutations at 29 positions, and 12 CTD mutations at 11 positions. CoV-RDB also contains results on 89 combinations of spike mutations in addition to VOCs and VOIs. These generally consist of sets of two or three mutations that are present within a VOC or VOI. Several of the most common mutation combinations are listed in the footnote of Table 2.

**Table 2.**
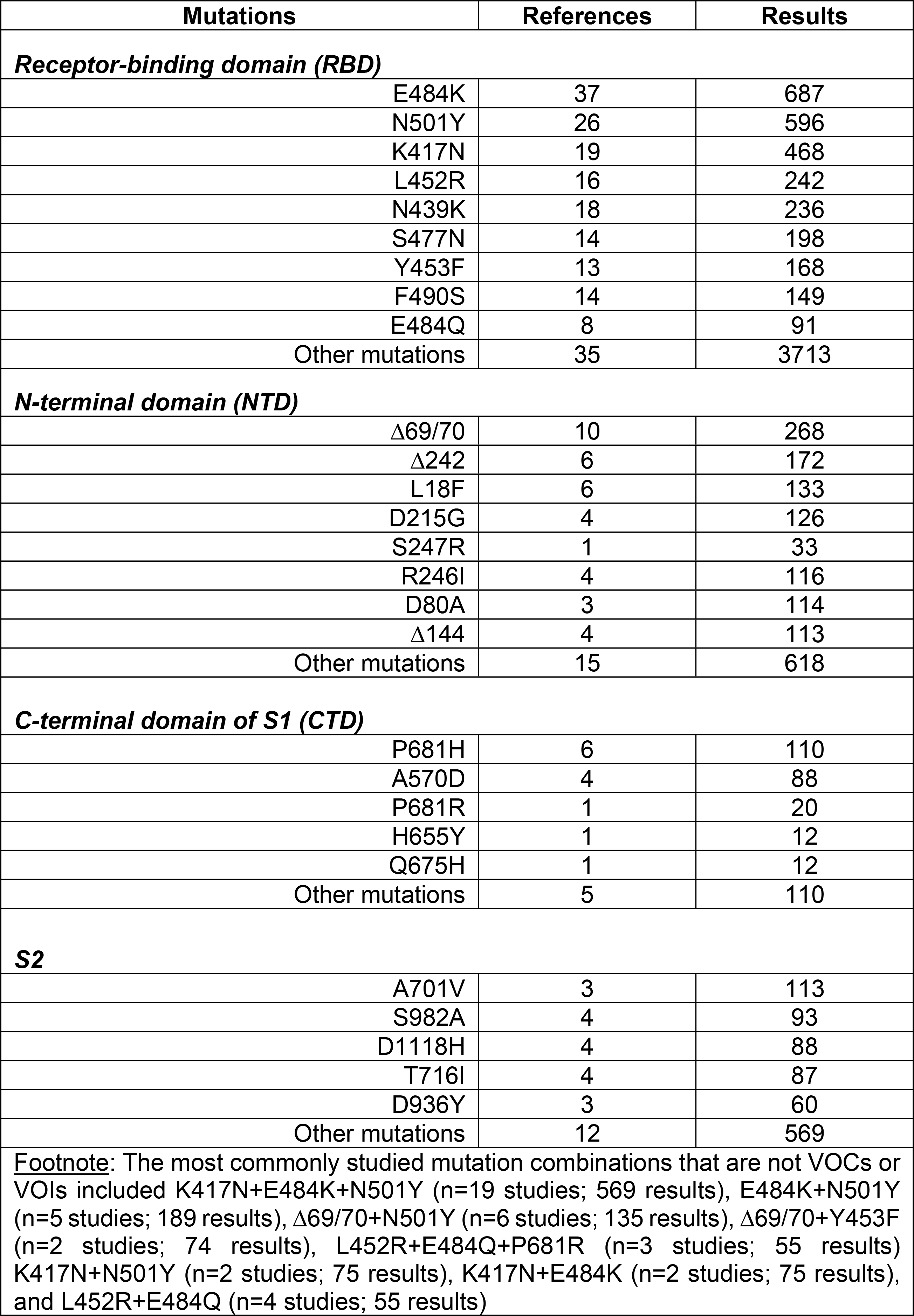
Individual SARS-CoV-2 Spike Mutations for which Data are Available in CoV-RDB

| **Mutations** | **References** | **Results** |
| --- | --- | --- |
| ***Receptor-binding domain (RBD)*** |  |  |
| E484K | 37 | 687 |
| N501Y | 26 | 596 |
| K417N | 19 | 468 |
| L452R | 16 | 242 |
| N439K | 18 | 236 |
| S477N | 14 | 198 |
| Y453F | 13 | 168 |
| F490S | 14 | 149 |
| E484Q | 8 | 91 |
| Other mutations | 35 | 3713 |
| ***N-terminal domain (NTD)*** |  |  |
| Δ69/70 | 10 | 268 |
| Δ242 | 6 | 172 |
| L18F | 6 | 133 |
| D215G | 4 | 126 |
| S247R | 1 | 33 |
| R246I | 4 | 116 |
| D80A | 3 | 114 |
| Δ144 | 4 | 113 |
| Other mutations | 15 | 618 |
| ***C-terminal domain of S1 (CTD)*** |  |  |
| P681H | 6 | 110 |
| A570D | 4 | 88 |
| P681R | 1 | 20 |
| H655Y | 1 | 12 |
| Q675H | 1 | 12 |
| Other mutations | 5 | 110 |
| ***S2*** |  |  |
| A701V | 3 | 113 |
| S982A | 4 | 93 |
| D1118H | 4 | 88 |
| T716I | 4 | 87 |
| D936Y | 3 | 60 |
| Other mutations | 12 | 569 |
| **Footnote:** The most commonly studied mutation combinations that are not VOCs or VOIs included K417N+E484K+N501Y (n=19 studies; 569 results), E484K+N501Y (n=5 studies; 189 results), Δ69/70+N501Y (n=6 studies; 135 results), Δ69/70+Y453F (n=2 studies; 74 results), L452R+E484Q+P681R (n=3 studies; 55 results) K417N+N501Y (n=2 studies; 75 results), K417N+E484K (n=2 studies; 75 results), and L452R+E484Q (n=4 studies; 55 results) |  |  |

### Antibodies (mAbs, CP, and VP)

#### mAbs

In CoV-RDB, mAbs are characterized according to their stage of clinical development; spike target; and specific epitope, i.e., the list of amino acids within 4.5 angstroms of the mAb paratope, according to structural data obtained from the Protein DataBank (PDB). Table 3 lists those mAbs with FDA emergency use authorizations (EUAs) or in advanced clinical development along with their spike targets and numbers of experimental results including *in vitro* selection and neutralization susceptibility data. In addition to the mAbs in Table 3, CoV-RDB contains 340 results for five mAbs in phase I/II trials; and 4975 results for 295 mAbs that are not currently in clinical development. For 54 of these 295, additional mAbs, structural data are available in the PDB. Table 4 summarizes neutralization susceptibilities for the six most studied variants and the six most studied individual spike mutations to those mAbs that either have EUAs or are in advanced clinical development.

**Table 3.**
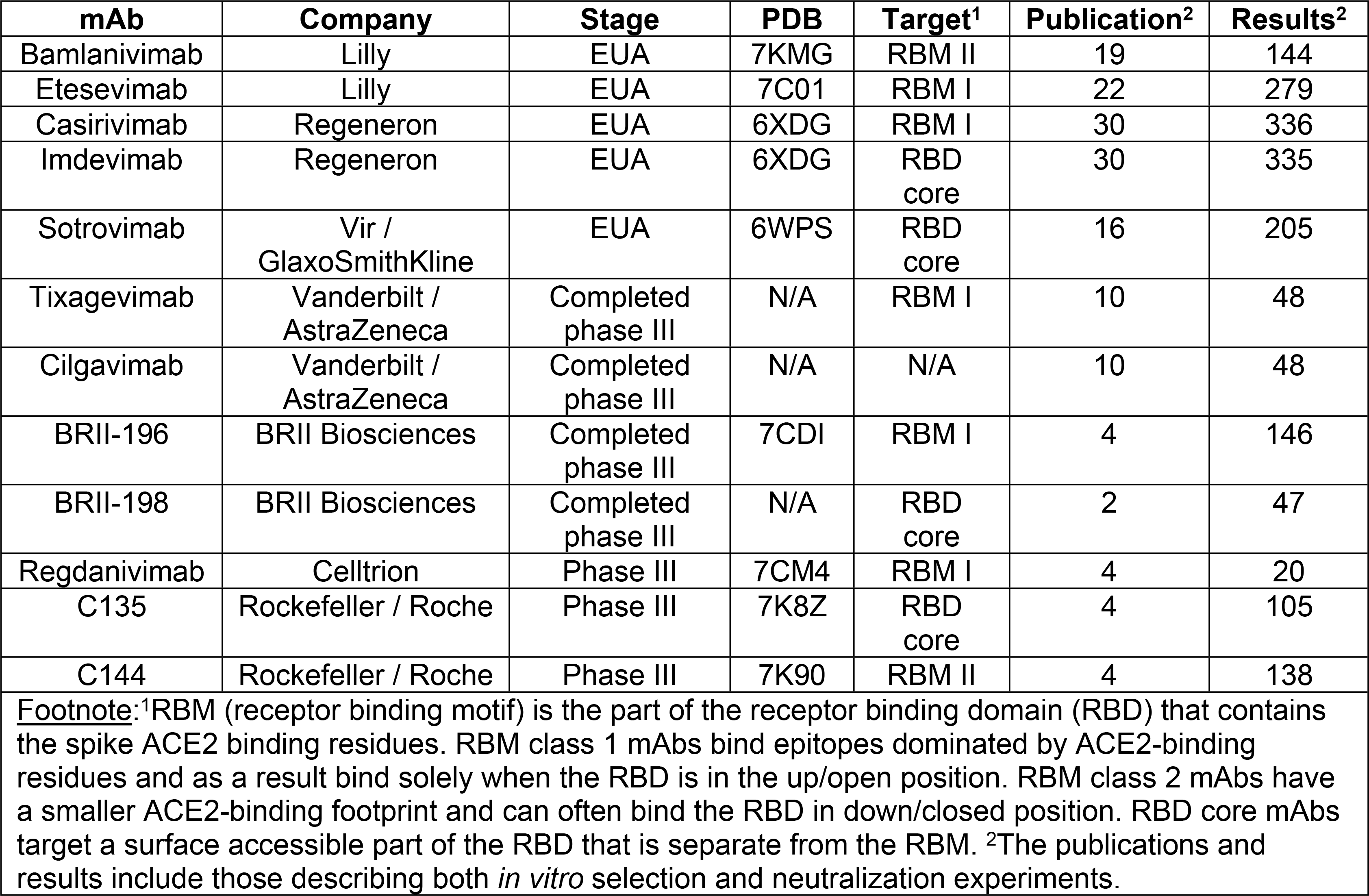
Monoclonal Antibodies (mAbs) with Emergency Use Authorization (EUA) or in Advanced Clinical Trials

**Table 4.**
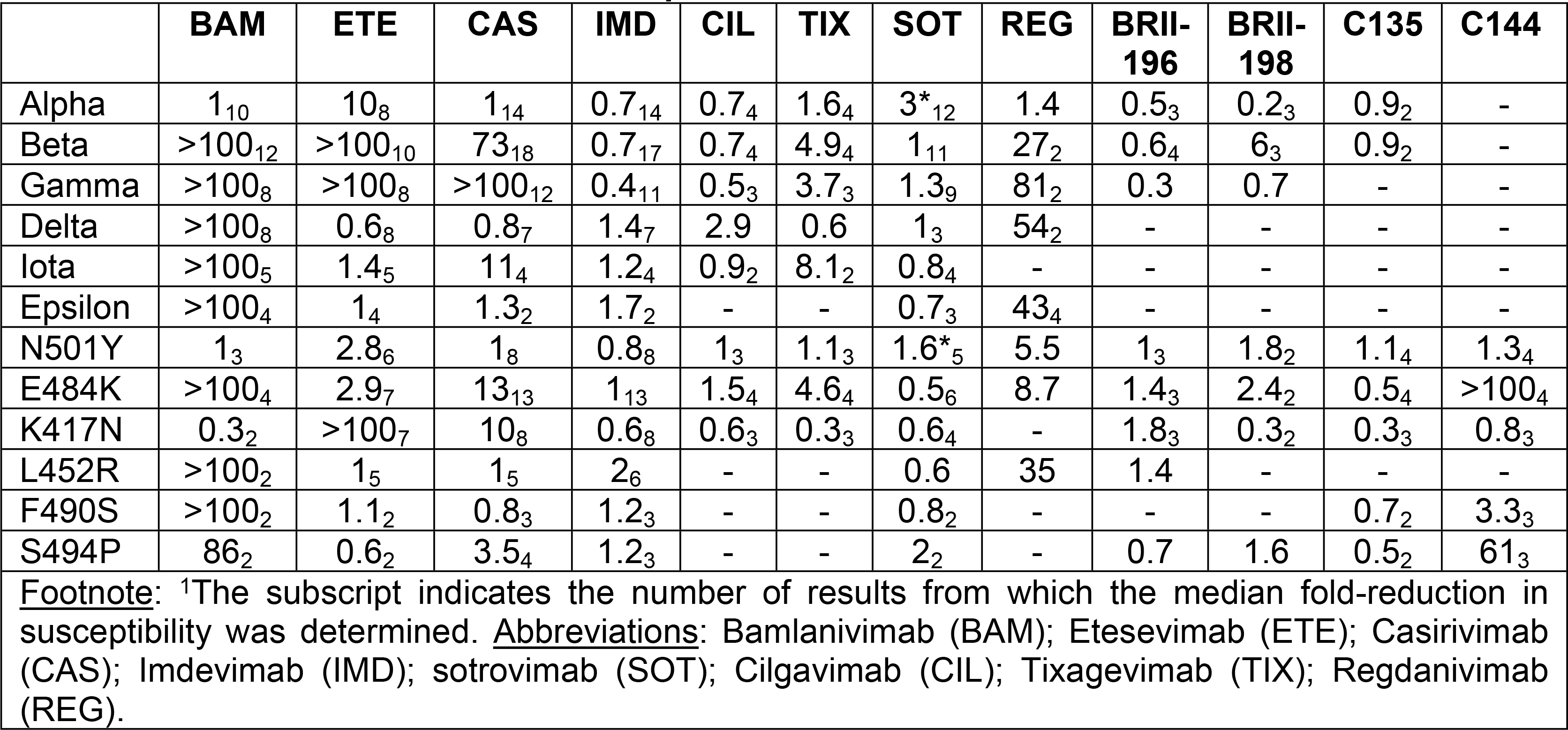
Median Fold Reduced Neutralization Susceptibility of 6 Variants and 6 Spike Mutations to 12 mAbs in Advanced Clinical Development^1^

Fig 1 shows the relationship between mAb epitopes, mutations selected during *in vitro* passage, and mutations that reduce mAb binding and/or susceptibility. S2 Fig shows that there is a strong correlation between RBD binding in a high throughput deep mutational scanning (DMS) assay and neutralization in that an escape fraction of >0.1 in the DMS assay is with few exceptions associated with a >10-fold reduction in neutralization in cell culture [8].

**Fig 1.**
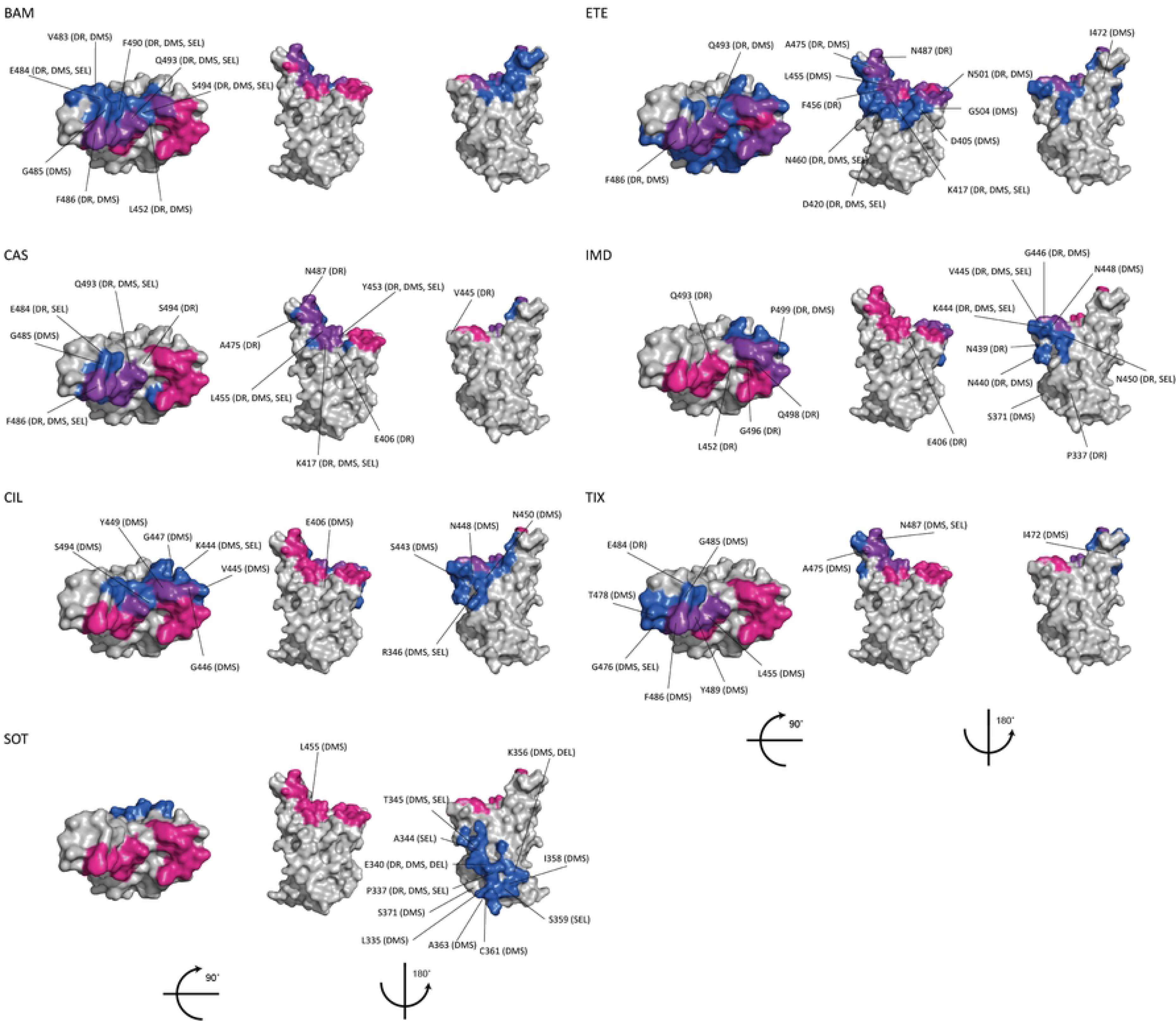
**Monoclonal antibodies (mAbs) with EUAs or in advanced clinical development: receptor binding domain (RBD) epitopes and immune escape positions**. For each mAb, the top of the RBD and two side views are depicted using coordinates from PDB 6M0J. ACE2 binding residues are shown in red; the mAb epitope defined as those residues within 4.5 angstroms of the RBD is shown in dark blue; and ACE2 binding residues within the mAb epitope are shown in purple. Those positions containing mutations that were either selected by the mAb *in vitro* (“SEL”), reduced binding in a deep mutational scanning assay (“DMS”), and/or reduced *in vitro* neutralizing susceptibility by a median of ≥4-fold in CoV-RDB (drug resistance; “DR”) are also indicated. The mAb epitopes for BAM (bamlanivimab), ETE (etesevimab), CAS (casirivimab), IMD (imdevimab), SOT (sotrovirmab), CIL (cilgavimab) and TIX (tixagevimab) were determined from their PDB structures.

#### CP

In CoV-RDB, CP are characterized by the sequence of the infecting variant, severity of illness, and time since infection. Table 5 lists the numbers of experimental results in CoV-RDB according to these characteristics and the SARS-CoV-2 variant or mutation(s) tested for neutralization. Overall, 11,390 neutralization experiments were performed using CP samples (96 studies) including 84 studies that provided data for individual samples and 12 studies that provided only aggregate data.

**Table 5.** Numbers of Convalescent Plasma (CP) Neutralization Experiments in CoV-RDB According to the Infecting Virus, Time Since Infection, and SARS-CoV-2 Variant Tested

| Characteristic | References | Results |
| --- | --- | --- |
| ***Infecting virus*** |  |  |
| Ancestral (A, B, B.1, others) | 86 | 8778 |
| Alpha | 10 | 825 |
| Beta | 7 | 214 |
| Gamma | 2 | 38 |
| Delta | 1 | 40 |
| ***Time since infection*** |  |  |
| 1 month | 74 | 6661 |
| 2-3 months | 35 | 2743 |
| 4-6 months | 23 | 1397 |
| >6 months | 14 | 589 |
| ***Variant / mutations undergoing neutralization*** |  |  |
| Alpha | 55 | 2294 |
| Beta | 58 | 2416 |
| Gamma | 26 | 964 |
| Delta | 13 | 444 |
| VOIs (kappa, iota, epsilon, lambda) | 9 | 161 |
| Other variants | 19 | 655 |
| Individual mutations and mutation combinations | 28 | 3795 |

The time since infection was available for all CP sample results including 10,801 samples obtained within six months of infection and 589 samples obtained beyond six months. The severity of illness was described for 3,559 (31%) CP samples with 2,133 samples from persons with mild-to-moderate disease and 1,426 samples from persons with severe disease. Although the sequence of the infecting virus was rarely known, the vast majority were obtained prior to the emergence of VOCs and VOIs. Nonetheless, 7%, 2%, 1%, and 1% were obtained from persons known to be infected with the Alpha, Beta, Gamma, and Delta variants, respectively.

#### VP

In CoV-RDB, VP are characterized according to the vaccine received, number of vaccinations, time since vaccination, and whether the VP was obtained from a person who was also previously infected with SARS-CoV-2. Table 6 lists the numbers of experimental results according to each of the above characteristics and the variant or mutation(s) tested for neutralization. 70% of VP were obtained from persons receiving one of the two widely used mRNA vaccines (BNT162b and mRNA-1273) while about 20% were obtained from persons receiving the Coronovac, AZD1222, or Ad26.COV2s vaccines. Approximately 80%, 18%, and 2% of VP samples were obtained within 1 month, 2-6 months, and >6 months after vaccination, respectively. Nearly 5% of samples were obtained from persons with confirmed infection prior to vaccination.

**Table 6.** Numbers of Vaccinee Plasma (VP) Neutralization Experiments in CoV-RDB According to the Vaccine, History of Previous Infection, Time Since Infection, and SARSCoV-2 Variant Tested.

| Characteristic | References | Results |
| --- | --- | --- |
| ***Vaccine*** |  |  |
| BNT162b2 | 82 | 10608 |
| mRNA-1273 | 31 | 4048 |
| CoronaVac | 8 | 1607 |
| AZD1222 | 15 | 1527 |
| Ad26.COV2.S | 8 | 828 |
| BBV152 | 5 | 449 |
| Sputnik V | 3 | 408 |
| AZD1222 + BNT162b2 | 3 | 258 |
| MVC-COV1901 | 1 | 342 |
| BBV154 | 1 | 120 |
| BBIBP-CorV | 2 | 111 |
| NVX-CoV2373 | 2 | 148 |
| AZD1222 + AZD2816 | 1 | 92 |
| ZF2001 | 2 | 56 |
| mRNA-1273 + mRNA-1273.351 | 1 | 40 |
| AZD2816 | 1 | 20 |
| ***Previously SARS-CoV-2 infected*** |  |  |
| Yes | 14 | 837 |
| No | 125 | 20035 |
| ***Time since vaccination*** |  |  |
| 1 month | 113 | 16789 |
| 2-3 months | 21 | 3072 |
| 4-6 months | 10 | 662 |
| > 6 months | 8 | 349 |
| ***Variant / mutations undergoing neutralization*** |  |  |
| Alpha | 73 | 4743 |
| Beta | 79 | 5090 |
| Gamma | 40 | 2114 |
| Delta | 32 | 2063 |
| VOIs (kappa, iota, epsilon, lambda) | 21 | 978 |
| Other variants | 38 | 1898 |
| Individual mutations and mutation combinations | 23 | 2327 |
| ***Number of vaccinations*** |  |  |
| 1 | 34 | 4438 |
| 2 | 118 | 15979 |
| 3 | 6 | 455 |
| **Footnote:** The numbers of references and results obtained from persons receiving uncommonly studied vaccines or vaccine combinations are not shown. |  |  |

Fig 2 shows the distribution of fold-reductions in neutralizing susceptibilities and absolute neutralizing antibody titers against each of the VOCs for previously uninfected persons completing the recommended course of vaccination for eight vaccines. A similar fold-reduction in neutralizing susceptibility against a given variant is likely to be more consequential for those vaccines that elicit lower titers of neutralizing antibodies. The figure shows that regardless of the vaccine, the fold reduction in neutralizing susceptibility was consistently greater for the Beta variant and that VP from persons receiving one of the two mRNA vaccines was usually more likely to retain neutralizing activity against each of the VOCs.

**Fig 2.**
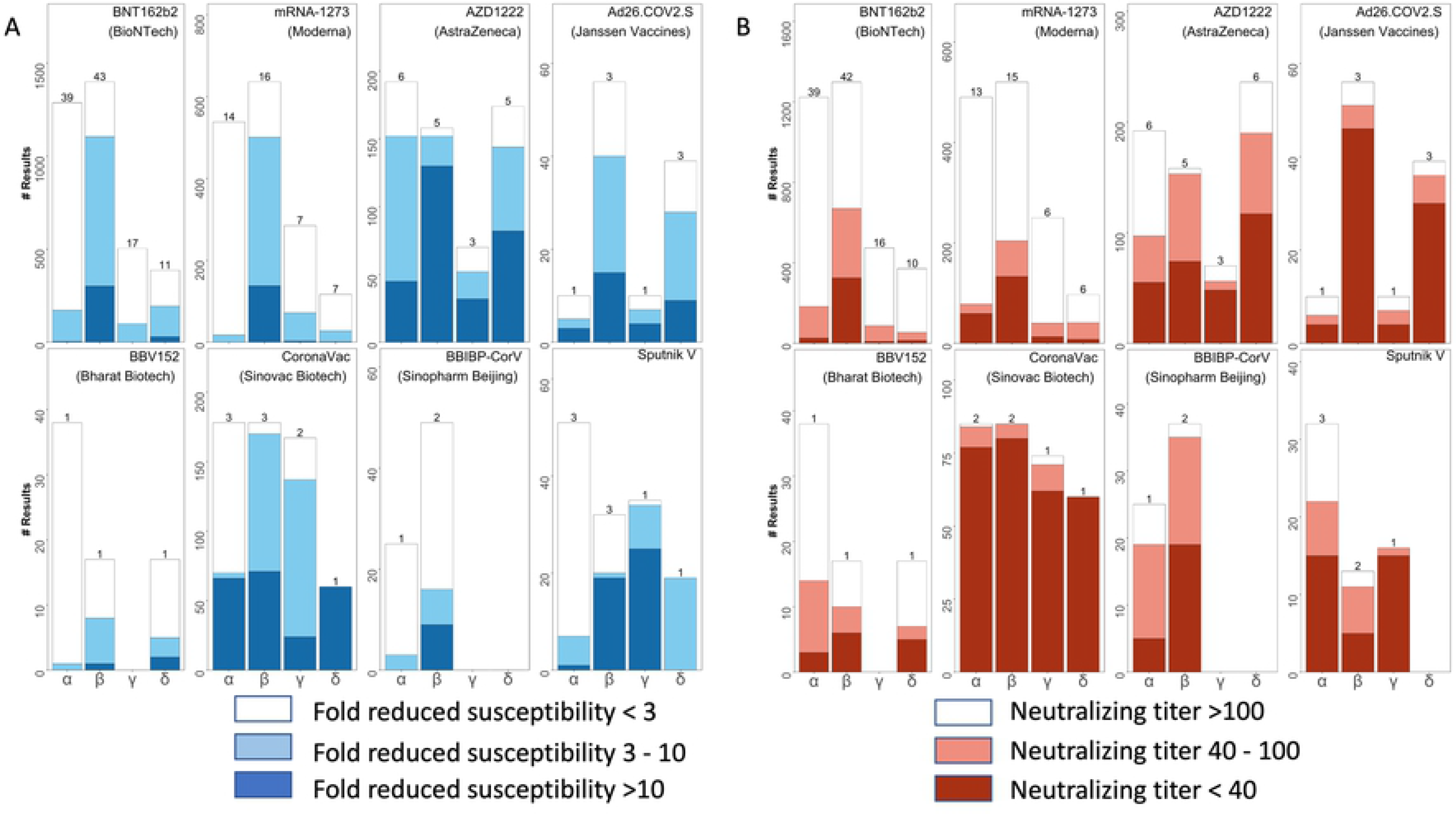
Distribution of fold-reduction in susceptibilities (A) and of absolute neutralizing titers (B) of vaccinee plasma (VP) associated with eight vaccines to the four variants of concern (VOC): Alpha, Beta, Gamma, and Delta. The X axes indicate the Greek letter associated with each VOC. The Y axes indicate the number of neutralizing assays. The numbers above the stacked bars indicate the number of studies reporting the experimental results. The figure includes VP obtained solely from previously uninfected persons receiving a complete vaccination schedule.

#### CoV-RDB Website

The CoV-RDB website contains four main features: (i) searchable tables containing SARS-CoV-2 variants, mAbs, vaccines, and references; (ii) regularly updated data summaries such as the data shown in Table 4 for mAbs and in Fig 2 for VP; (iii) user-defined queries; and (iv) a sequence analysis program. The user-defined queries and sequence analysis program are described here because they are interactive and contain multiple user options.

### User-Defined queries

#### Query interface

The query interface allows users to search the database using one or more of the following three criteria: (i) published reference; (ii) antibody preparation (mAb, CP, or VP); and (iii) SARS-CoV-2 variant or spike mutation(s). If the “References” dropdown option is selected, then all the data associated with that reference is displayed in separate tables containing neutralization susceptibility data for mAbs, CP, and VP and/or *in vitro* selection data for mAbs. If the “Plasma / mAbs” dropdown option is selected, users must select either a specific mAb, CP from a person infected with a specific variant, or VP from persons who received a specific vaccine. If the “Variants / Mutations” dropdown option is selected, users must select a particular VOC, VOI, other variant, individual spike mutation, or combination of spike mutations. Selecting items from multiple dropdown boxes restricts the output to the data specified by the combination of dropdown items. Fig 3 shows the output returned by selecting E484K from the “Variants / Mutations” dropdown. Fig 4 shows the output returned by selecting BNT162b from the “Plasma / mAbs” dropdown and the Delta variant from the “Variant / Mutations” dropdown.

**Fig 3.**
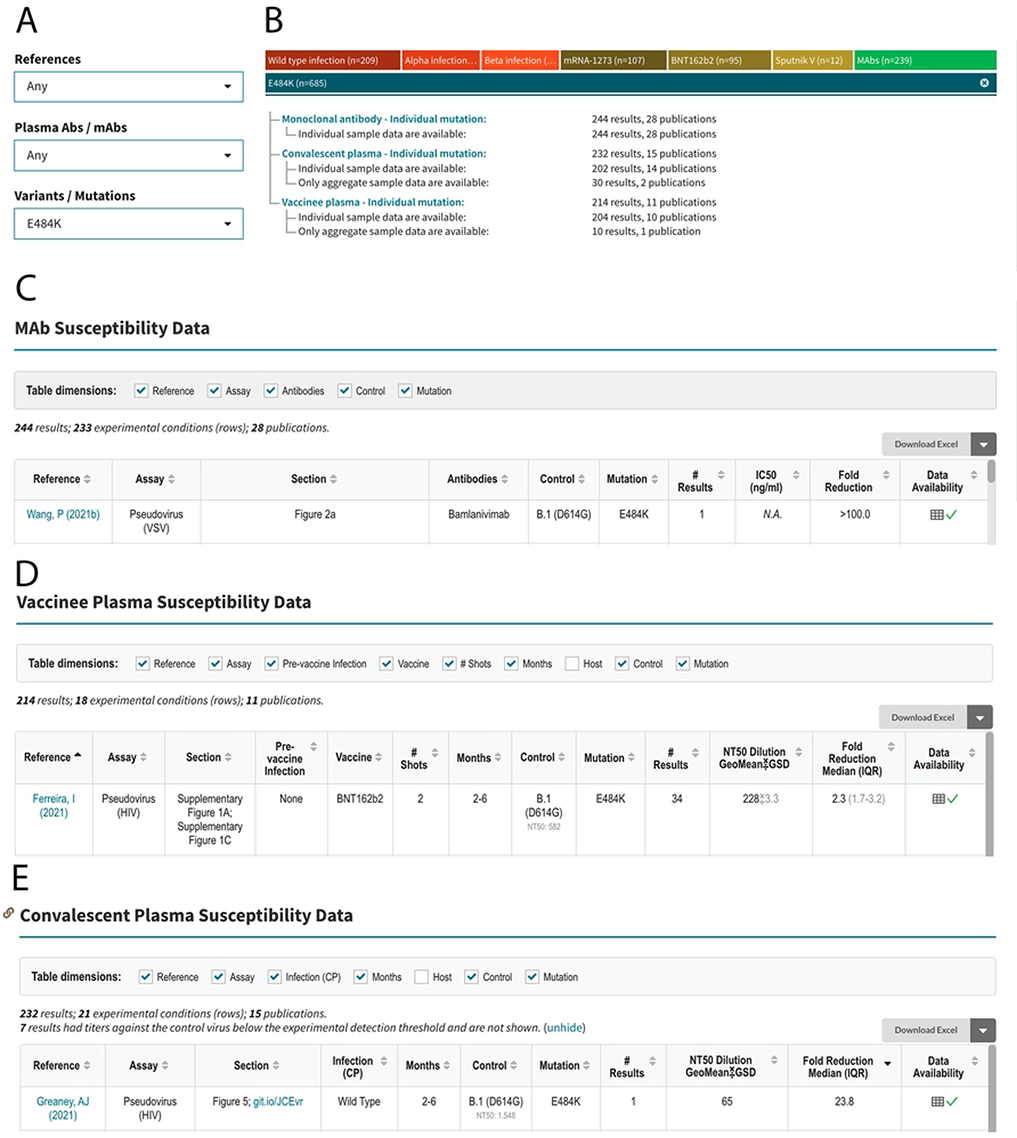
Query interface, sample query, and outline of query results for an example query for the SARS-CoV-2 RBD mutation E484K. The query interface containing three dropdown boxes is shown at the upper left (A). E484K is selected from the “Variants / Mutations” dropdown box. The upper right summarizes the data returned by the query, which in this case includes 244 mAb neutralizing susceptibility results from 28 publications, 232 convalescent plasma (CP) results from 15 publications, and 214 vaccinee plasma (VP) results from 11 publications (B). The summary distinguishes between results for which only aggregate (mean or median) data are provided and results for which individual data are provided. The sections below show the headers and first row of the tables containing mAb (C), VP (D), and CP (E) susceptibility results. The complete contents of these tables can be found on the web (https://covdb.stanford.edu/search-drdb/?host=human&mutations=S%3A484K).

**Fig 4.**
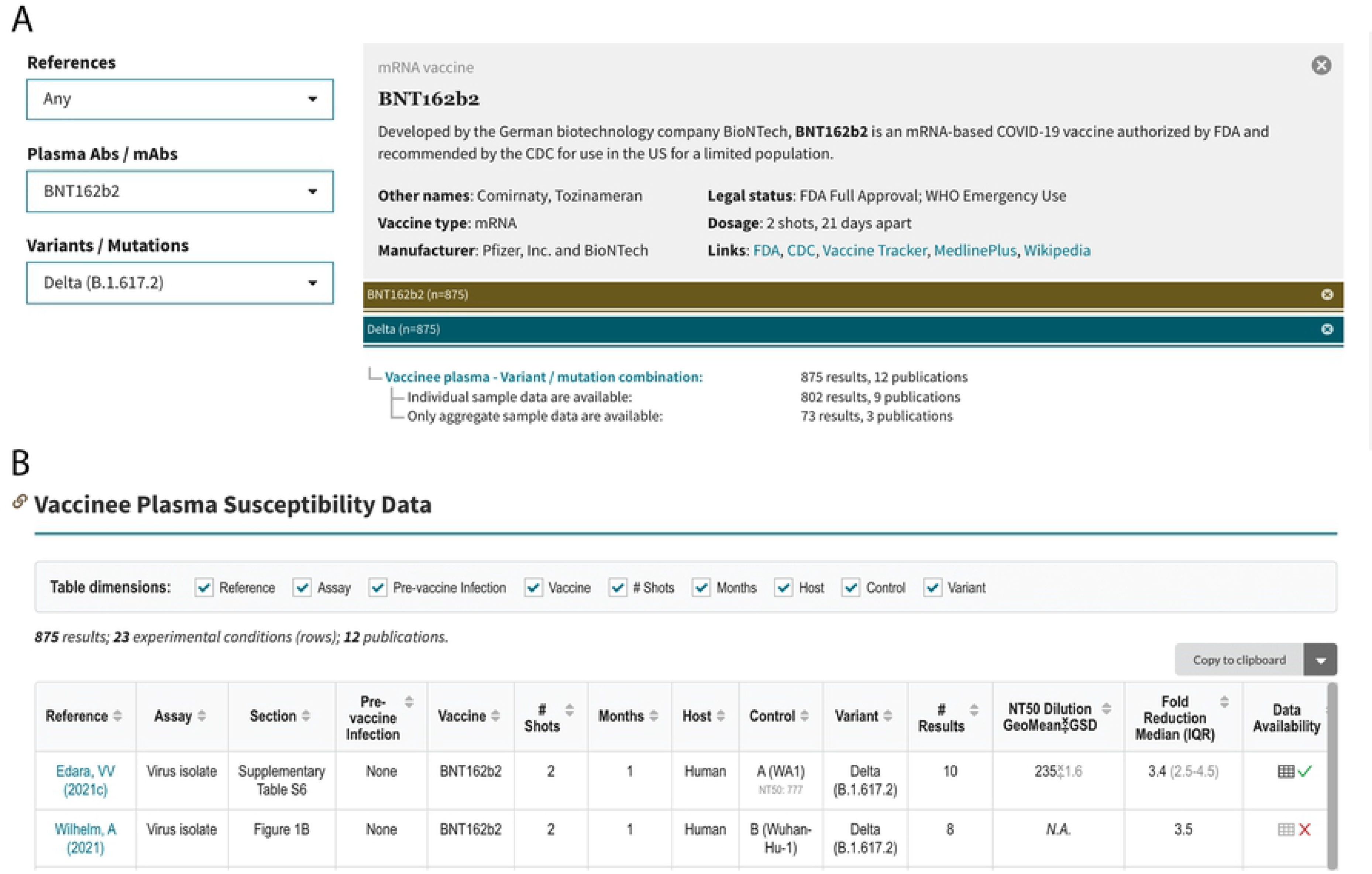
Query interface, sample query, and outline of query results for an example query for the susceptibility of the SARS-CoV-2 Delta variant to the vaccinee plasma (VP) of persons who received the BNT162b vaccine. The query interface containing three dropdown boxes is shown at the upper left (A). BNT162b is selected from the “Plasma / mAbs” dropdown box, and the Delta variant is selected from the “Variants / Mutations” dropdown box. The upper right summarizes the data returned by the query, which in this case includes 875 results from 12 publications (B). The summary distinguishes between results for which only aggregate (mean or median) data are provided and results for which individual data are provided. The section below the header shows the header and first few rows of the table entitled “Vaccinee Plasma Susceptibility Data”. The complete contents of this tables can be found on the web (https://covdb.stanford.edu/search-drdb/?host=human&vaccine=BNT162b2&variant=Delta).

#### Query results

The mAb, CP, and VP susceptibility query result tables contain between 10 to 13 column headers (Fig 3C-3E). The mAb table contains column headers indicating the reference and location of the data within each reference (e.g., figure or table); assay type (e.g., pseudotyped virus); mAb tested; variant tested; IC_50_ in ng/ml; and fold-reduced susceptibility compared with a control virus (that is also present in the table; Fig 3C).

The VP table contains column headers that indicate the reference and location of the data within the reference, type of assay used, vaccine received, number of immunizations, number of months since immunization, whether the sample was obtained from a vaccinated person with confirmed prior infection, variant tested, geometric mean neutralizing titer, and median fold reduction in titer compared with the control virus (Fig 3D and Fig 4C).

The CP table contains column headers that indicate the reference and location of the data within the reference, assay type, lineage of the virus that infected the person from whom the CP was obtained, number of months between infection and the plasma sample, variant tested, geometric mean neutralizing titer; and median fold reduction in titer compared with the control virus (Fig 3E).

Each table also contains two additional columns: “# Results” and “Data Availability”. The number of results indicates the number of neutralizing experiments. The data availability contains a “√” if individual data are available or an “X” if data are only available in aggregate (i.e., as a geometric mean titer or a median fold reduction in susceptibility). Clicking on the spreadsheet icon in “Data Availability” column copies the contents of that study to the user’s clipboard. The “Download CSV” tab at the top of each table allows users to download all rows for analysis.

#### Interacting with the query results

Each table can be considered to have multiple dimensions represented by the table’s column headers. For example, Fig 5A indicates the table dimensions for the query output with BNT162b selected from the “Plasma Abs / mAb” dropdown and the Delta variant selected from the “Variants / Mutations” dropdown. The table contains 22 rows (i.e., experimental conditions) summarizing 885 results obtained from 13 references. The table dimension check boxes enable users to aggregate the data in the table by deselecting one or more dimensions. For example, if the user deselects the “Reference”, “Assay”, and “Control” variant dimensions, the data are summarized with six rows that provide the median fold-reduction in susceptibility to BNT162b-associated VP for the Delta variant according to the number of vaccinations received, the time since vaccination, and whether the VP was obtained from a previously vaccinated person (Fig 5B).

**Fig 5.**
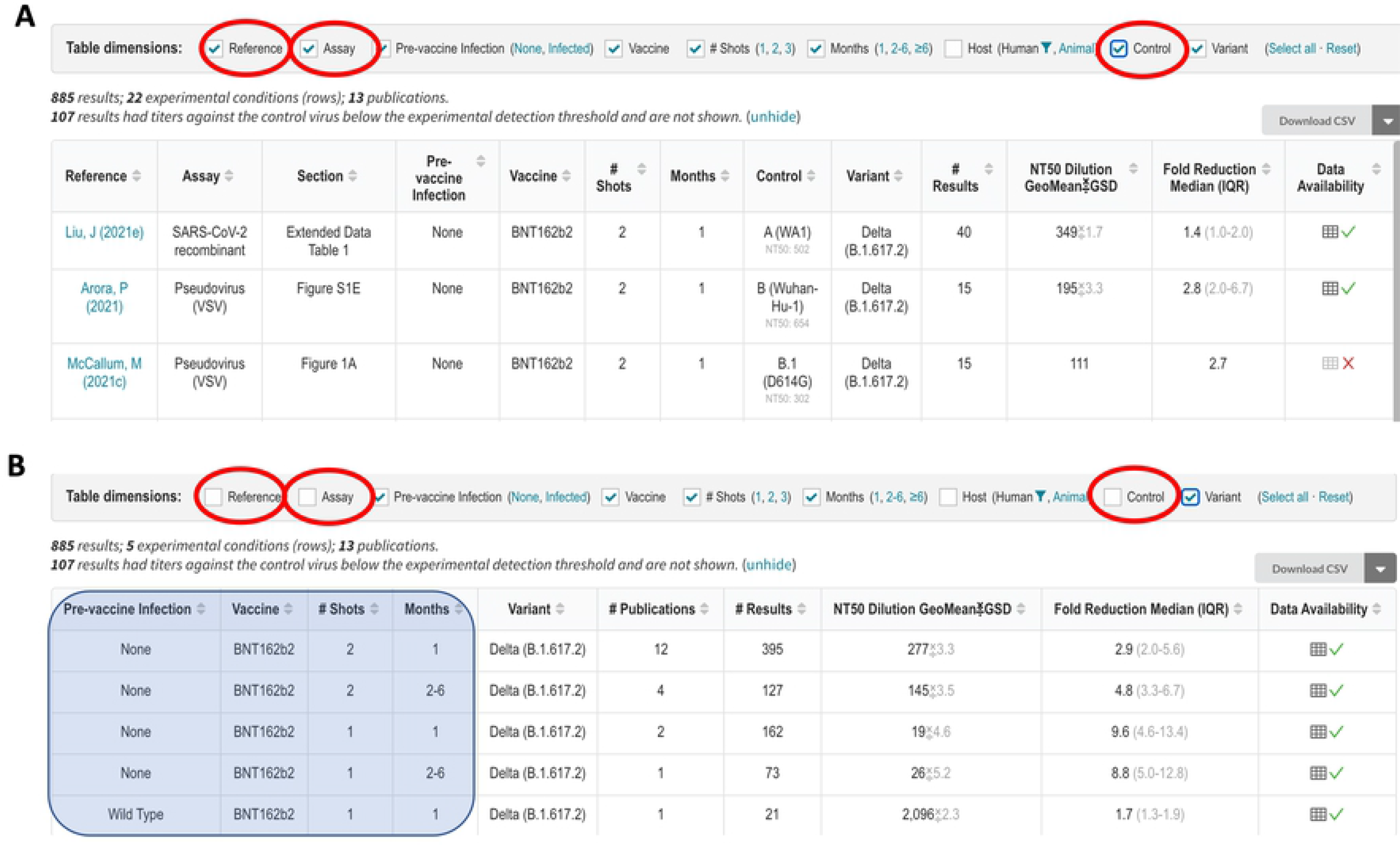
Aggregating the query results for the example shown in. **Fig 4**. The top part of the figure shows the first three of 22 experimental conditions defined by the column headers in which BNT162b-associated VP was tested for activity against the Delta variant (A). The results for each experimental condition are shown in different rows each containing the geometric mean of the neutralizing antibody titer and the median fold-reduction in titer compared with a control virus. The bottom part of the figure shows that the 22 rows can be displayed using 6 rows by aggregating those results obtained from different references using different assays or different control viruses by deselecting the “Reference”, “Assay”, and “Control” variant (checkboxes within red ovals). The neutralization data are now aggregated according to the number of BNT162b immunizations, time since immunization, and whether the VP was obtained from a person who had been infected prior to vaccination (shaded area superimposed on the first four columns). The number of experimental results in each row in Fig 5B is increased because each row may now contain data from more than one reference.

### Sequence analysis program

The CoV-RDB website contains a SARS-CoV-2 sequence analysis program that leverages the same code base written for the widely used Stanford HIV Drug Resistance Database interpretation program [9, 10]. The CoV-RDB sequence analysis program supports three types of input: (i) a list of spike mutations; (ii) one or more consensus FASTA sequences containing any part of the SARS-CoV-2 genome; and (iii) one or more codon frequency table (CodFreq) files containing the following seven columns: gene, amino acid position, number of reads at that position, codon, amino acid encoded by the codon, number of reads for that codon, and proportion of reads for that codon. An auxiliary program provided through the website or via download enables users to convert next generation sequencing (NGS) FASTQ files into human and machine-readable CodFreq files that are much faster to analyze. CodFreq files make it possible to estimate the extent of background noise resulting from sequencing or PCR artifact (S3 Fig) [10].

If one or more spike mutations is submitted, the CoV-RDB sequence analysis program reports information on specific mutations and generates summary tables containing the susceptibility of viruses with these mutations to mAbs, CP, and VP (S4 Fig). If a FASTA sequence is submitted, the program returns comments, summary tables, a list of the SARS-CoV-2 genes in the submitted sequence, a list of each of the amino acid mutations in each virus gene, and the sequence’s PANGO lineage (S4 Fig). If a codon frequency table is submitted, the program returns all of the above information and also summarizes the read coverage for each position along the genome and provides users with options to select read depth and mutation detection thresholds below which mutations will not be reported (S4 Fig). If multiple sequences are submitted, users have the option of obtaining all the results in a single downloadable CSV file.

Fig 6A-6C show three parts of the output generated when either a FASTQ sequence or CodFreq file is submitted to the sequence analysis program: sequence summary (Fig 6A), sequence quality assessment (Fig 6B), and mutation list (Fig 6C). The mutation comments and neutralization susceptibility data, which are also included in the output, are not shown. The sequence summary section (Fig 6A) lists the genes that underwent sequencing, the median read depth, and the Pango lineage. This section also contains three dropdown boxes that help users select the appropriate threshold for identifying sub-consensus mutations. The read depth and mutation detection thresholds are used to select the minimum number and proportion of reads required for a mutation to be considered viral in origin rather than a sequence or PCR artifact. The nucleotide mixture threshold allows users to select a threshold which minimizes the number of nucleotide ambiguities present in the sequence.

**Fig 6.**
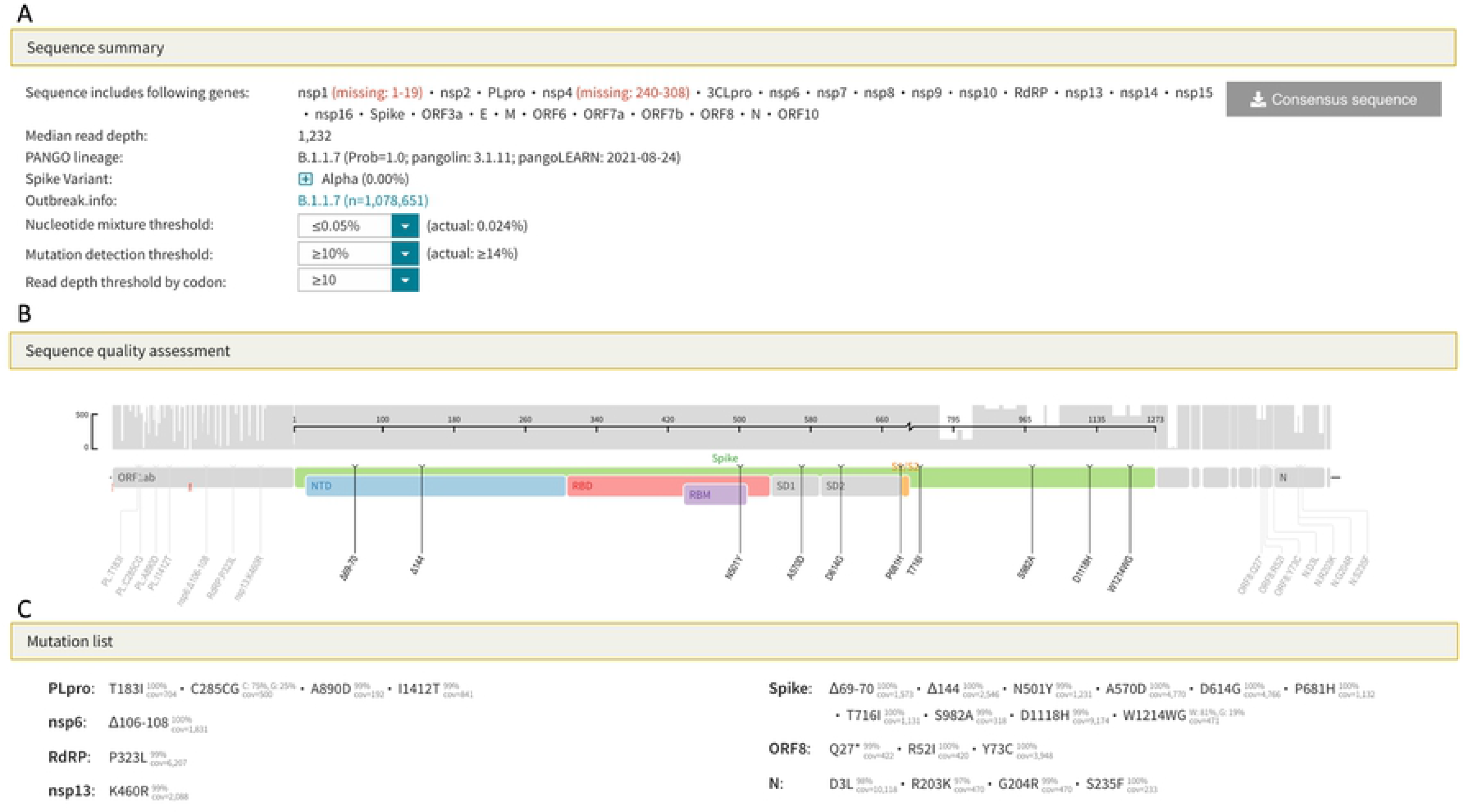
SARS-CoV-2 sequence analysis program output for FASTQ sequences or codon frequency tables. The sequence summary section (A) lists the genes that were sequenced, the median read depth, and the Pango lineage. This section also contains dropdown boxes that enable the user to select the minimum number and proportion of reads, respectively, required to identify a mutation. The threshold that minimizes the proportion of positions with nucleotide ambiguities can also be selected. The sequence quality assessment section displays the read depth across the genome and lists only those amino acid mutations that meet the user-defined criteria outlined in the sequence summary section (B). The mutation list section lists those amino acid mutations that meet the user-defined criteria and shows the proportion of reads containing the mutation (C). The output shown in this figure can be regenerated by loading the example file B.1.1.7 (ERR5026962) at this URL: https://covdb.stanford.edu/sierra/sars2/by-reads/.

## Discussion

Because SARS-CoV-2 is a rapidly evolving RNA virus infecting millions of people worldwide, new variants will continue to emerge and influence regional and global pandemic trajectories. Ongoing epidemiological and clinical studies are needed to understand the risk of re-infection and vaccine failure posed by different variants. However, because the spectrum of SARS-CoV-2 variants is expanding and shifting faster than epidemiological studies can be conducted, laboratory-based markers will increasingly be used to identify those variants that could predispose to re-infection and vaccine failure. SARS-CoV-2 neutralizing antibody titers are correlates of immune protection and are likely to be useful as an endpoint in vaccine trials and to determine the need for vaccine boosters and immunogen updates.

CoV-RDB is the only database to comprehensively curate published data on the neutralizing susceptibility of SARS-CoV-2 variants and spike mutations to mAbs, CP, and VP. Although non-neutralizing Abs and cellular immune responses also contribute to protection from infection, the presence of neutralizing antibodies targeting the spike protein has correlated most strongly with protection from infection in animal models and in previously infected and vaccinated persons [11–14], Neutralizing susceptibility assays are also better suited to standardization compared with assays of other immunological defense mechanisms.

CoV-RDB can be downloaded in its entirety without restrictions. This is accomplished using a dual database pipeline that combines the full-fledged Postgres database system to enforce relational data integrity and the simplicity of the SQLite database system to enable users to download and query the database without the overhead of accessing a host server. By making the database fully available to all users, we aim to encourage data sharing and the editing of the underlying CSV files by the authors of published studies.

Cov-RDB neutralizing susceptibility query output can be considered multidimensional tables in which the rightmost columns contain numerical results (e.g., titers and fold-reductions in susceptibility) while the leftmost columns contain experimental conditions. The experimental conditions are explanatory variables that either directly influence neutralizing susceptibility (e.g., specific vaccine, time since vaccination, and SARS-CoV-2 variant) or have a more subtle effect on susceptibility (e.g., type of neutralizing assay and control virus). The CoV-RDB query interface enables users to explore the query results at different levels of granularity by filtering or aggregating them according to one or more experimental conditions without making additional calls to the web server.

The sequence analysis program shares many features with the Sierra HIV Drug Resistance Database sequence analysis program. For the analysis of NGS data, both programs use Minimap2 [15] to align individual reads to a reference sequence resulting in SAM/BAM files and SAM2CodFreq, a program we wrote to create CodFreq files. The CodFreq format has several advantages over the commonly used variant call format (VCF) because it can be interpreted without a reference sequence and can be used independently from the accompanying SAM file. CodFreq files have a simple tabular format enabling them to be viewed and manipulated using a spreadsheet. The sequence analysis program uses the CodFreq files to assist users determine the appropriate threshold for distinguishing background sequence artifact from authentic sub-consensus amino acids.

### Study limitations

Neutralizing susceptibility data are highly heterogeneous often resulting in discordant results across studies [16, 17]. There are three main sources for this heterogeneity. First, the composition of neutralizing antibodies among previously infected and vaccinated individuals is heterogeneous [6, 18]. Second, there are different types of neutralizing assays including those performed in cell culture using pseudotyped viruses, chimeric viruses, recombinant viruses, and clinical isolates and, more recently, surrogate neutralizing assays that assess the ability of antibodies to block the interaction between the SARS-CoV-2 RBD and ACE2. Third, results for the same sample against a virus variant can differ even among laboratories using the same type of assay as a result of differences in virus inoculum size, the cells used for culture, and viral replication endpoints [16,17,19]. As neutralizing assays become more standardized and as external controls such as those provided by the WHO [20] are increasingly used, it is likely that reproducibility across studies will improve.

Data on the clinical significance of SARS-CoV-2 neutralizing susceptibility is continually evolving [11,21–26]. The utility of neutralizing antibodies as a correlate of immune protection is ultimately determined in epidemiological studies. Therefore, a database devoted to protective immunity should ideally contain both laboratory and epidemiological data. The main obstacle to expanding CoV-RDB to also include epidemiological studies of vaccine efficacy is that such studies are much more complex than those reporting *in vitro* neutralizing data. For example, vaccine efficacy data depends not just on the vaccine, the variant, and the time since vaccination but also on the study design and the age and immune status of the study population. Moreover, in many vaccine efficacy studies, the proportion of individuals infected with different variants is not known.

### Future directions

Although the CoV-RDB is centered around just four main entities (references, viruses, antibodies, and experiments), differences among the types of viruses, types of antibody preparations, and types of experiments has necessitated a sophisticated database design. Nonetheless, as new types of experiments are being published, we have been expanding the database schema. For example, an increasing number of studies report the results of surrogate neutralizing assays based on antibody-mediated blocking of the interaction of RBD with ACE2 [19,27,28]. These assays appear to correlate strongly with plasmid and conventional neutralization test results, although they do not assess the effects of NTD-binding and other non-RBD antibodies that synergistically inhibit virus replication [19]. In addition, the use of an international external standard for calibration such as the one developed by the WHO will increase concordance across different assays and will be reported using international units rather than as an IC_50_ (for mAbs) or a plasma dilution [20, 29].

We have also added the comprehensive deep mutational scanning data published by the Bloom laboratory to the database [8,30–34]. The data from these studies are displayed only for those mAbs in advanced clinical development and those mutations that have been reported to occur at a frequency above 0.001%. In addition to reporting the escape fraction associated with a mutation to an mAb, we report the level of protein expression within yeast of RBDs containing the mutation (a measure of protein stability) and the ACE2 binding of RBDs containing the mutation as mutations that bind poorly to ACE2 are less likely to be selected *in vivo*. Although binding data have been reported for many other mAbs using enzyme linked immunoassays, surface plasmon resonance, and biolayer interferometry, we have not curated these data as they have not been as comprehensive as the deep mutational scanning data.

Commercial total binding assays do not differentiate between binding and neutralizing antibodies. They also do not measure binding or neutralization of multiple variants but rather assess binding to pre-variant spike proteins. However, total binding assays often display moderately strong correlations with neutralizing assays [29,35,36] and activity against specific variants may eventually be assessed using variant-specific reagents. We may eventually add such data to CoV-RDB if they will provide insights that cannot be obtained solely from neutralizing antibody studies.

Although CoV-RDB contains neutralizing susceptibility data obtained using the plasma from infected animal experiments (e.g., non-human primates, hamsters, and mice), we have not included data from animal model challenge studies as such studies would require extensive modifications to our current database schema. Therefore, we will continue to monitor these studies and consider adding top-line data from these studies to alert database users to the existence of these studies without rigorously representing study details. Finally, a similar approach will be considered for studies of vaccine efficacy. We are therefore exploring the possibility of adding the top-line data of these studies so that the findings of these studies can be correlated with the *in vitro* neutralizing data in CoV-RDB.

## Supporting information

**S1 Table.** Slightly different patterns of spike mutations within each of the SARS-CoV-2 variants of concern (VOCs) and variants of interest (VOIs).

**S1 Fig. Data management workflow for CoV-RDB.** Weekly incremental searches of PubMed and preprint servers (BioRxiv/MedRxiv and Research Square) are performed. Publications that appear to have data pertinent to SARS-CoV-2 variants and their susceptibility to mAbs, convalescent plasma (CP), and vaccinee plasma (VP) are downloaded to a Zotero reference database folder to enable full-text review and data curation. Extracted data are exported into a set of linked CSV files in an open-source Github repository (https://github.com/hivdb/covid-drdb-payload). Extracted data are then imported into a Postgres database where the data are validated for completeness and consistency before being exported as a single SQLite database file that serves as the back end for the CoV-RDB website and is available to users for download.

**S2 Fig.** Effects of spike receptor binding domain (RBD) mutations on monoclonal antibody (mAb) binding and neutralization. The X-axis shows the escape fraction as determined using a deep mutational scanning (DMS) assay that measures the binding of various mAbs to RBD variants produced in yeast. The Y-axis shows the results of in vitro neutralization using the same mAbs and spike proteins with the same RBD mutations as those used in the DMS assay. Discordant results are shown in the figure. Abbreviations: SOT: sotrovimab; BAM: bamlanivimab; ETE: etesevimab; CAS: casirivimab; IMD: imdevimab.

**S3 Fig. Creating a codon frequency file from a FASTQ file.** FASTQ files are aligned to the consensus Wuhan reference sequence using the Minimap2 alignment program. The resulting BAM/SAM files are then processed by a program SAM2CodFreq that we wrote to generate a codon frequency file containing seven columns as shown on the right. The table here shows the results from three codons (spike positions 500 to 502). The observation that many codons shown in this (and other parts of the same file which are not shown) are present at levels between 0.2% and about 2% suggests that codons present at these low proportions likely represent sequencing or PCR artifact (i.e., “background noise”). However, as the mutation N501Y occurs at a considerably higher proportion (34.3%), it is likely to be present in the infecting virus population.

**S4 Fig. Functions of the SARS-CoV-2 sequence analysis program.** The program supports three types of input: a list of spike mutations; one or more consensus FASTA sequences containing any part of the SARS-CoV-2 genome; and one or more FASTQ sequences. However, because a FASTQ sequence can take several minutes to analyze, users are advised to first convert them to a codon frequency file through an auxiliary program. If a list of spike mutations is submitted, the program returns comments about notable mutations and summary tables reporting the susceptibility of viruses with these mutations to mAbs, CP, and VP. If a FASTA sequence is submitted, the program returns the preceding information plus a list of the SARS-CoV-2 genes, the amino acid mutations in the sequence, and the sequence’s PANGO lineage. If a FASTQ sequence or codon frequency table is submitted, the program provides the preceding information and the read coverage for each position along the genome. It also provides users with the options to select read depth and mutation detection thresholds below which mutations will not be reported.

